# Genome-scale CRISPR screening reveals host factors required for ribosome formation and viral replication

**DOI:** 10.1101/2022.11.12.515717

**Authors:** Maikke B. Ohlson, Ashwani Kumar, Jennifer L. Eitson, Seoyeon Jang, Chunyang Ni, Chao Xing, Michael Buszczak, John W. Schoggins

## Abstract

Viruses are known to co-opt host machinery for translation initiation, but less is known about which host factors are required for the formation of ribosomes used to synthesize viral proteins. Using a loss of function CRISPR screen, we show that synthesis of a flavivirus encoded reporter depends on multiple host factors, including several 60S ribosome biogenesis proteins. Viral phenotyping revealed that two of these factors, SBDS, a known ribosome biogenesis factor, and the relatively uncharacterized protein SPATA5, were broadly required for replication of flaviviruses, coronaviruses, alphaviruses, paramyxoviruses, an enterovirus, and a poxvirus. Mechanistic studies revealed that loss of SPATA5 caused defects in ribosomal RNA processing and ribosome assembly, suggesting that this human protein may be a functional ortholog of yeast *Drg1*. These studies implicate specific ribosome biogenesis proteins as viral host dependency factors that are required for synthesis of virally encoded protein and accordingly, optimal viral replication.

## INTRODUCTION

As obligate parasites, viruses depend on host cellular machinery for nearly every step of their replication cycles. Advances in genome-wide screening technologies enabled by HAP1 cells, siRNA gene-silencing, and CRISPR/Cas9 gene editing have elevated our knowledge of the host factor repertoire required by viruses. In many cases, these screens have identified host factors that are uniquely required for specific viruses. For example, CRISPR enabled identification of virus-specific entry factors CD300lf for norovirus (Orchard et al., 2016) and the alphavirus receptor Mxra8 (Zhang et al., 2018). Additionally, comparative screening performed with dengue, West Nile, yellow fever, hepatitis C, and Zika virus infections identified protein processing complexes required across the *Flaviviridae*, including the Sec61 translocon, the oligosaccharyltransferase (OST) complex, and ER membrane complex (EMC) (Krishnan et al., 2008), (Marceau et al., 2016), (Lin et al., 2017), (Zhang et al., 2016), (Savidis et al., 2016), (Ma et al., 2015), (Barrows et al., 2019), (Hoffmann et al., 2021).

In addition to virus family-specific host factors, host proteins that are more broadly utilized by diverse classes of viruses have also been identified through genetic screens. Examples include molecules involved in the vacuolar ATPase, heparan sulfate biosynthesis, endocytic trafficking, the COG complex, and translation initiation, among others (Sessions et al., 2009), (Perreira et al., 2015), (Hoffmann et al., 2017), (Tanaka et al., 2017), (Riblett et al., 2016), (Li et al., 2020).

Notably, host factor screens have not identified many proteins that regulate translation outside of the initiation steps. These genes may have been missed because the screen relied on death-based selection strategies, which counter-select against essential host genes, or used siRNA strategies that fail to achieve complete gene silencing. Because coopting translation machinery is a hallmark of all viruses, we hypothesized that additional host factors beyond those already identified may be important for viral protein expression, and by extension, genome replication.

Here, we used a genetically tractable flavivirus, yellow fever virus expressing the Venus green fluorescent protein, to implement a genome-scale CRISPR knockout screen that relied on virally encoded reporter gene expression as the selection strategy, rather than cell survival after virus-induced cell killing. This approach revealed numerous known flavivirus host factors as well as a cluster of 60S ribosome biogenesis factors that were not identified in previous screens. SBDS, a known ribosome biogenesis factor, and the relatively uncharacterized SPATA5 were validated in targeted knockout studies. We further demonstrate that these factors are broadly required for infection by multiple viruses representing 12 RNA and DNA viral families. Loss of each host factor affects polyribosome formation, thereby blunting replicative potential. SPATA5 was demonstrated to have a role in ribosomal RNA processing, implicating it as a functional ortholog to yeast *Drg1*. Together, these data identify previously uncharacterized viral host dependency factors and suggest a model in which specific proteins, some of which regulate ribosome biogenesis, are required to achieve a critical threshold of viral protein production required for optimal replication.

## RESULTS

The goal of this study was to implement a CRISPR screen to identify host factors required for virally encoded protein synthesis during a single replication cycle. Prior to the screen, we first interrogated the effects of single cycle flavivirus infection on host ribosomes by infecting Huh7.5 cells with either the 17D strain of a yellow fever virus (YFV-17D) or a similar Venus-expressing virus (YFV-17D-Venus). Lysates were fractionated after sucrose gradient centrifugation, which separates the small subunit (40S), the large subunit (60S), individual monosomes (80S) and mRNAs with multiple ribosomes attached (polysomes) based on sedimentation rates. With infection, the level of polysomes was reduced and we observed a concomitant increase in 80S monosomes when compared to uninfected cells (Figure 1A), suggesting a defect in host mRNA translation elongation. An additional increase in the 60S subunit peak, but not the 40S subunit peak, was observed with both viruses. Similar profiles have been observed in Huh7 cells infected with dengue virus (DENV) and West Nile virus (WNV) (Roth et al., 2017).

**Figure 1.**
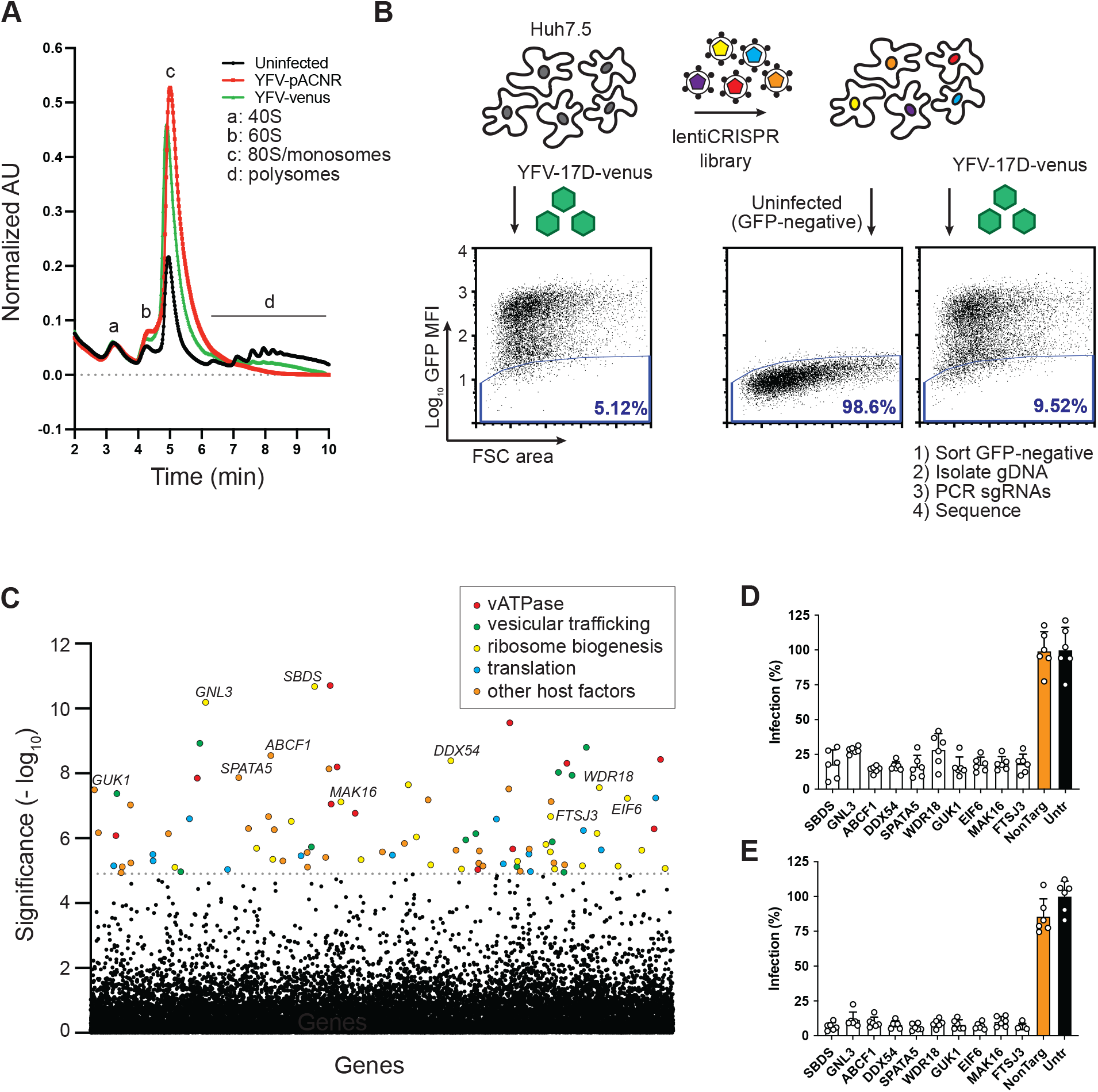
Yellow fever virus host factor CRISPR screen identifies 60S biogenesis host factors. A. Sucrose-density gradient ribosome fractionation profile of uninfected Huh7.5 cells (black), or infected with MOI 5 YFV-17D (red) or YFV-Venus (green) viruses for 24 h. B. Schematic of genome-wide “negative-selection” CRISPR screen to identify host factors required for YFV replication, blue line indicates FACS gate for GFP-negative cells. C. Manhattan dot plot of CRISPR screen results with significance of enrichment calculated by MAGeCK. Genes with FDR<0.01 (dotted line) are colored. D-E. Infectivity of Huh7.5 cells treated with CRISPR gRNAs targeting candidate genes infected with MOI 1 YFV-Venus for 24hr (D) or MOI 0.1 YFV-Venus for 48hr (E) and analyzed by FACS. Abbreviations: “NonTarg”, non-targeting CRISPR guides; “Untr”, untransduced control.

We hypothesized that flaviviruses may have a unique requirement for host factors that modulate some aspect of ribosome biology. To identify these putative host factors, we used a CRISPR-Cas9 “negative selection” strategy to enrich for sgRNA-expressing cells that failed to produce virally encoded Venus when infected with YFV-17D-Venus. In non-CRISPR treated Huh7.5 cells, YFV-17D-Venus infected approximately 95% of cells, with 5% of cells remaining Venus-negative (Figure 1B). However, in Huh7.5 cells transduced with the human CRISPR Brunello library (Sanson et al., 2018), the percentage of GFP-negative cells during YFV-17D-Venus infection increased to ~9.5%, likely representing an enrichment of cells lacking genes required for any viral infection step up to and including virally encoded Venus expression. The Venus-negative cell population was collected by FACS, and significantly enriched sgRNAs were identified by MAGeCK. (Li et al., 2014) (Figure 1C and Table S1). Analysis of significantly enriched genes (FDR < 1 %) using STRING database (Szklarczyk et al., 2021) revealed multiple pathways that have previously been identified in flavivirus host factor screens (Le Sommer et al., 2012), (Marceau *et al*., 2016), (Savidis *et al*., 2016), (Krishnan *et al*., 2008), (Lin *et al*., 2017), (Zhang *et al*., 2016), (Sessions *et al*., 2009), (Ma *et al*., 2015), (Li et al., 2019b), (Hoffmann *et al*., 2021), (Scaturro et al., 2018), (Shah et al., 2018), (Li et al., 2019a), (Barrows *et al*., 2019). These include genes regulating the vATPase complex, ER-Golgi/vesicular trafficking, translation initiation, and heparan sulfate biosynthesis (Figure S1A). We also identified 19 genes that mapped to a 60S ribosome biogenesis cluster that had not been identified in previous flavivirus studies, suggesting an important role for the 60S subunit in the YFV replication cycle.

To identify genes unique to our screen, we compared the <25% FDR candidate list to gene lists from flavivirus host factor screens performed using siRNA and CRISPR (Krishnan *et al*., 2008), (Sessions *et al*., 2009), (Le Sommer *et al*., 2012), (Ma *et al*., 2015), (Savidis *et al*., 2016), (Marceau *et al*., 2016), (Lin *et al*., 2017), (Li *et al*., 2019b), (Barrows *et al*., 2019), (Wang et al., 2020). We selected for validation 10 high-scoring genes that had not been previously identified in other viral host-factor screens: SBDS, GNL3, ABCF1, DDX54, SPATA5, WDR18, GUK1, EIF6, MAK16, and FTSJ3. To confirm the role of each gene in supporting YFV replication, two distinct Brunello library sgRNAs targeting each gene candidate were cloned into either a puromycin- or a blasticidin-selectable lentiCRISPRv2 vector. Dual knock-out (KO) Huh7.5 cells were generated by transducing cells with both lentiviruses and selecting cells in the presence of puromycin and blasticidin for 9 days to ensure high efficiency gene silencing and protein depletion. When candidate KO and control cells were infected with a high multiplicity of infection (MOI) of YFV-17D-Venus for 24hr, or approximately one replication cycle, we observed a potent reduction in the percentage of infected KO cells when compared to control cells (Figure 1D). Similar results were obtained at low MOI and 48 h (Figure 1E), confirming the requirement for these factors in the YFV replication cycle. We also tested how loss of several of these host factors affected YFV infectivity in two other human cell lines. We observed reduced infection with ABCF1, SPATA5, and GUK1 knockout, whereas loss of SBDS has less of an effect on YFV in these cells (Figure S1B, S1C).

Seven of the ten factors selected for follow up were annotated to be 60S ribosome biogenesis factors, and therefore the loss of each one may have contributed to impairment of viral infection via a common 60S subunit-based pathway. Since SBDS functions as the final co-factor in 60S maturation by removing EIF6 from the pre-60S particle (Warren 2018), we chose it for further analysis as the representative 60S ribosomal biogenesis factor out of the seven validated. We also chose the uncharacterized protein SPATA5, as it contains two ATPase domains and resembles yeast Afg2/Drg1, which regulates yeast 60S biogenesis factor recycling (Pertschy et al., 2007).

To determine whether loss of SBDS or SPATA5 directly contributed to impaired viral infection, we reconstituted CRISPR-targeted cells with cDNAs expressing CRISPR-resistant versions of each host factor gene, or Firefly luciferase (Fluc) as a control. KO cells expressing Fluc control were unable to support high levels of YFV-17D-venus infection, whereas KO cells that were complemented with each CRISPR-resistant host factor were as permissive to infection as control cells. (Figure 2A). We used western blot to confirm both CRISPR-mediated loss and cDNA-mediated reconstitution of each host factor (Figure 2B). These results indicate that SBDS and SPATA5 are each specifically required for efficient YFV infection.

**Figure 2.**
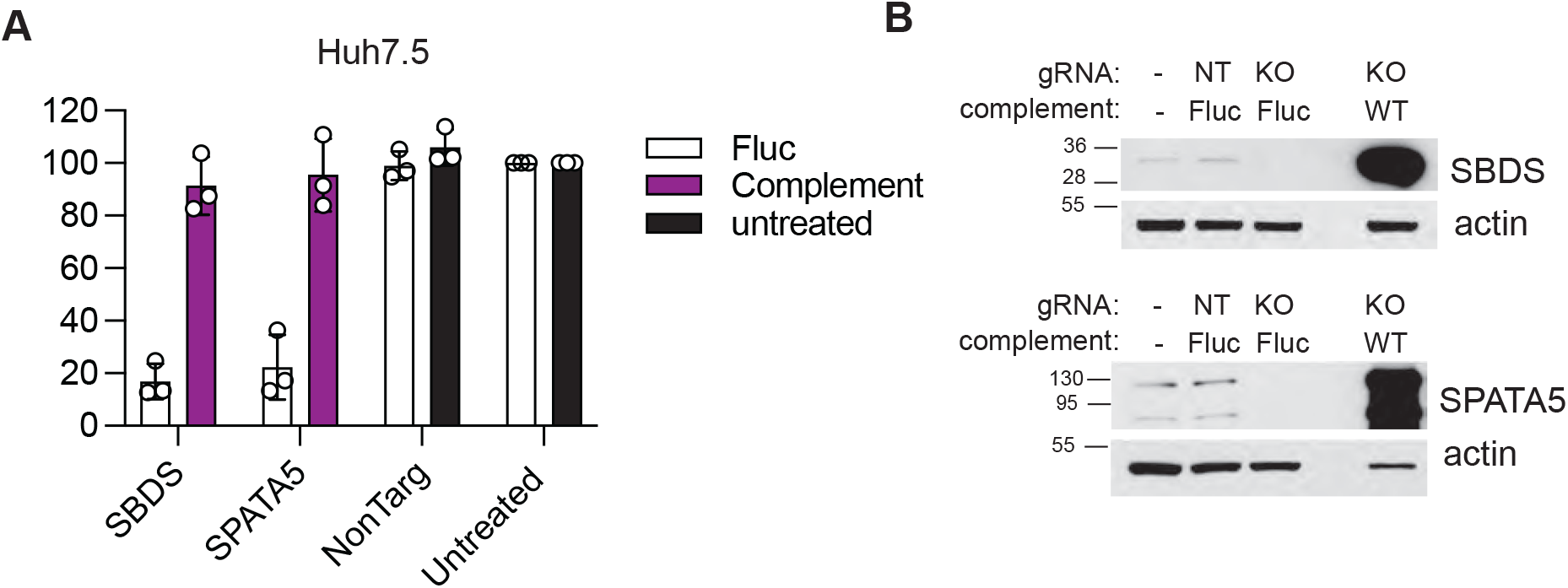
Genetic reconstitution of viral infection by expression of SBDS and SPATA5. A. Infectivity of Huh7.5 cells treated with indicated CRISPR gRNAs and complemented with lentiviral-expression of Fluc or guide-resistant host factors infected with MOI 1 YFV-Venus for 24hr and analyzed by FACS. B. Lysates from Huh7.5 cells untreated or treated with indicated CRISPR gRNAs and complemented with lentiviral expression of Fluc or guide-resistant host factors were analyzed by western blot using anti-SBDS or anti-SPATA5 antibodies. Blot membranes were also probed with anti-actin antibodies (indicated). Abbreviations: “NonTarg” or “NT”, non-targeting CRISPR guides; “Untr”, untransduced control.

Since virally encoded Venus is a surrogate reporter for viral replication, we asked whether loss of SBDS or SPATA5 affected infectious viral particle production of the parental non-reporter virus YFV-17D. Control and KO Huh7.5 cells were infected with YFV-17D at 0.01 MOI, and virus production was quantified over time by plaque assay (Figure 3A). Loss of either SBDS or SPATA5 resulted in log-fold reductions in YFV-17D titers. Similar results were obtained for the flaviviruses Zika virus (ZIKV) and dengue virus (DENV) (Figure 3B-C), suggesting a broader requirement for these host factors in the *Flaviviridae* family.

**Figure 3.**
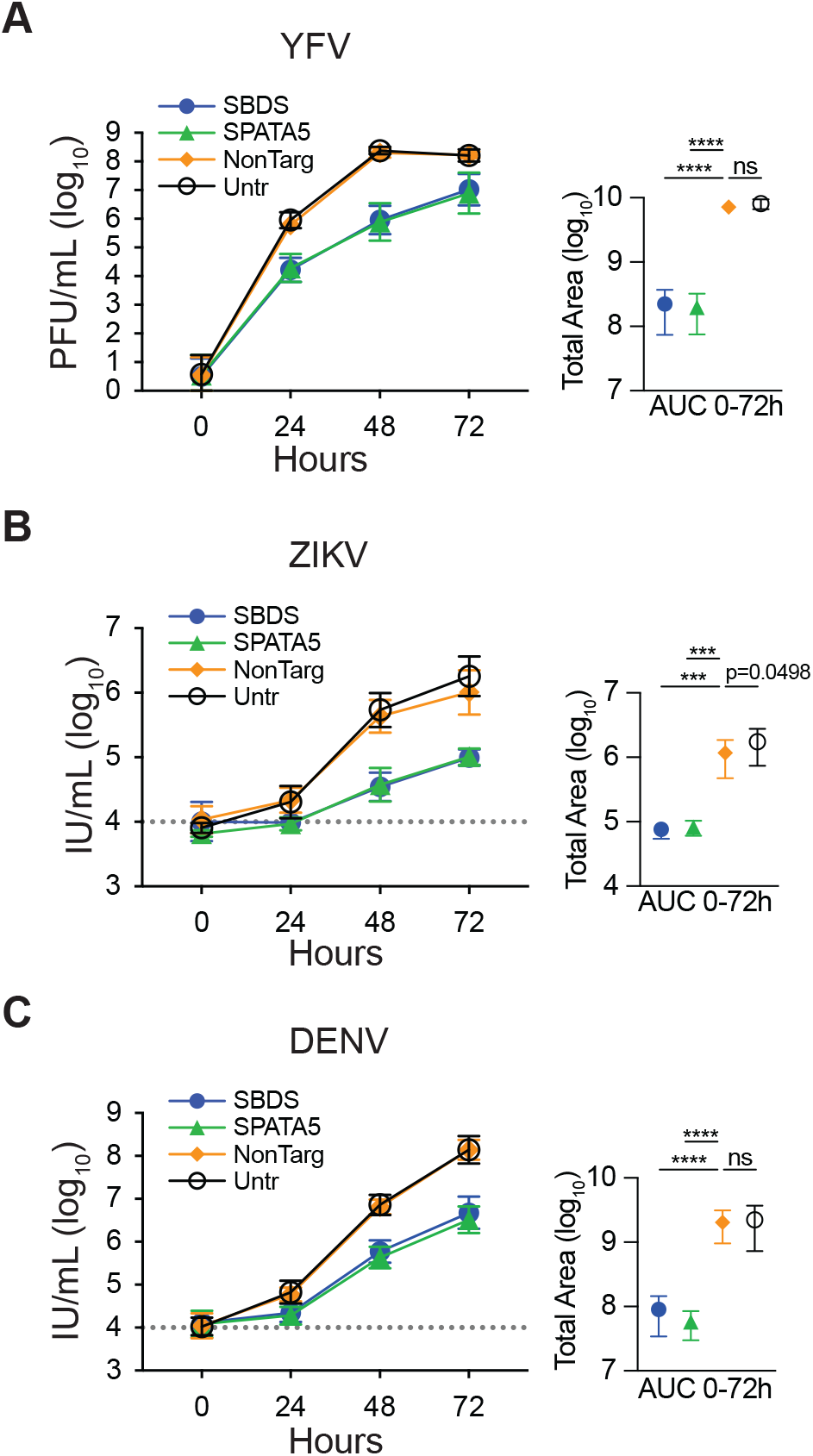
Flavivirus viral production is dependent on SBDS and SPATA5. A-C. (Left panels) Viral growth curves from control or KO Huh7.5 cells infected with 0.01 MOI of YFV-17D quantified by plaque assay (A), 0.1 MOI ZIKV PRVABC159 (B) or 0.1 MOI DENV2 (C). ZIKV and DENV2 titers determined by reinfection on susceptible Huh7.5 cells and antigen-staining by FACS (Right panels, dotted line indicates limit of detection). Area under the curve (AUC) was calculated and used to test statistical significance of each condition compared to NonTarg by one-way ANOVA. n=4, ***, P<0.001;; ****, P<0.0001

We next tested whether loss of SBDS or SPATA5 directly impaired synthesis of virally encoded proteins. For these assays, we used a replication-defective HCV replicon, which is a naked subgenomic RNA in which viral structural genes have been replaced with a secreted Gaussia luciferase (Gluc) reporter (Figure 4A). The RNA-dependent RNA polymerase is also mutated (GDDàGNN) so that the Gluc signal is detected only from translation of the transfected RNA. In contrast, a wild-type replicon self-amplifies, leading to logarithmic increases in Gluc signal over time (Figure 4B). When the GNN replicon was transfected into SBDS or SPATA5 knockout cells, we observed decreased Gluc synthesis at all time points (Figure 4C). Area under the curve analysis indicated that total Gluc levels were significantly reduced in the KO cells relative to non-targeting control. The differences between Gluc levels in control and KO cells were amplified considerably (2-3log_10_) with the WT replicon at later stages of replication (Figure 4D). This suggests that small defects in pioneering rounds of translation have pronounced consequences with respect to viral replication and virus production.

**Figure 4.**
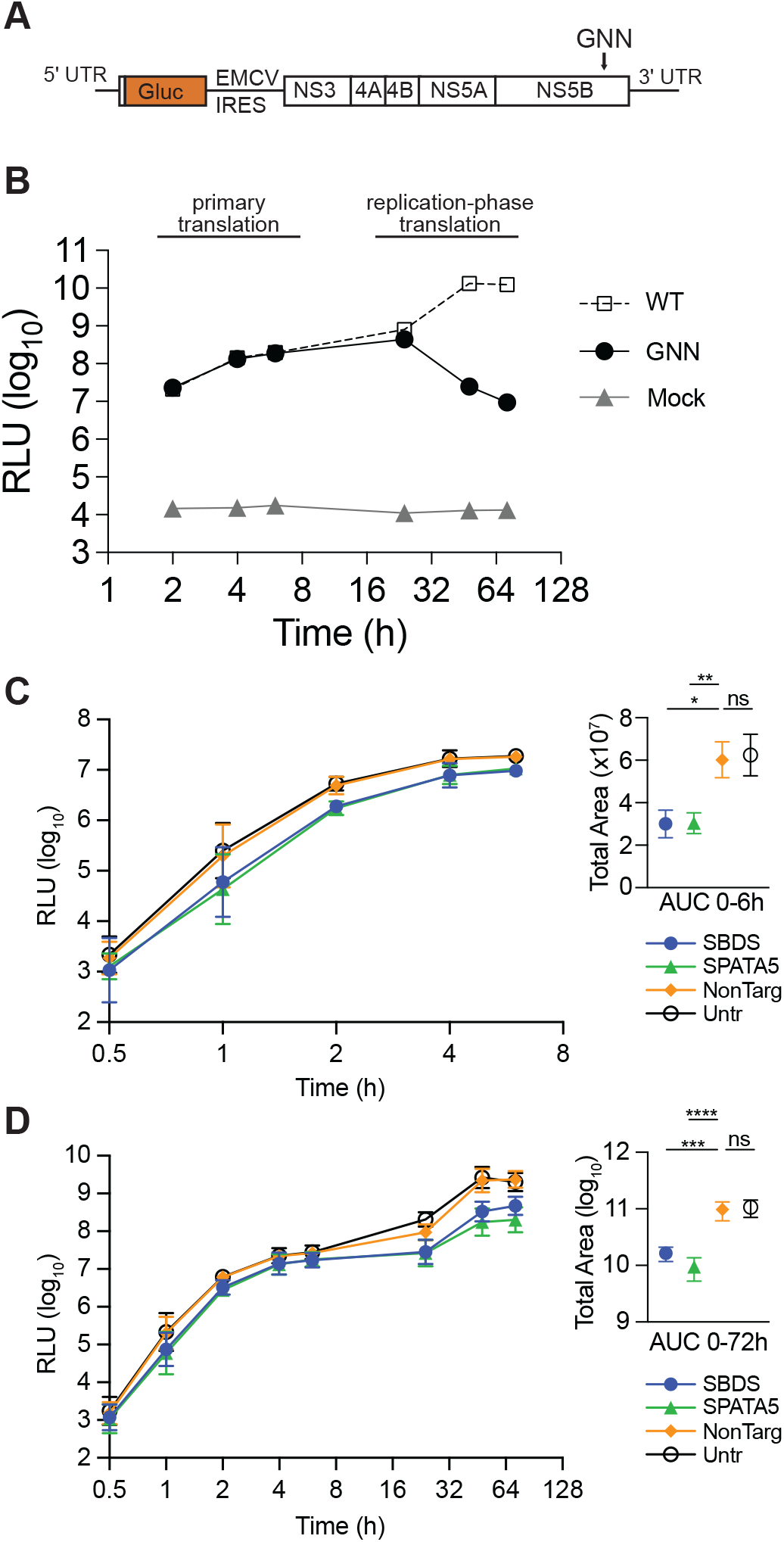
Virally encoded protein synthesis is reduced in SBDS and SPATA5 KO cells. A. Cartoon of HCV subgenomic replicon. B. Gluc synthesis from a replication-defective (GNN) and replication competent (WT) HCV subgenomic replicon in Huh7.5 cells. C-D. (Left panels) Gluc synthesis from replication-defective “GNN” (C) or replication-competent “WT” (D) HCV subgenomic replicon in KO and control Huh7.5 cells. (Right panels) Area under the curve (AUC) was calculated and used to test statistical significance of each condition compared to NonTarg by one-way ANOVA. n=4, *, P<0.05; **, P<0.01; ***, P<0.001; ****, P<0.0001. Abbreviations: “NonTarg”, non-targeting CRISPR guides; “Untr”, untransduced control.

We next tested whether the dependency of viruses on SBDS and SPATA5 extended beyond the *Flaviviridae*. We infected KO and control Huh7.5 cells with a panel of 15 diverse viruses, representing several negative- and positive-sense single-stranded RNA viruses, a double-stranded RNA virus, and double-stranded DNA viruses (Table S2). Viral infectivity was quantified by flow cytometry, using virally encoded GFP or virus-specific antigen staining. We observed that each host factor was required for optimal infection of cells by coxsackie virus B (CVB), equine arteritis virus (EAV), Sindbis virus (SINV), Venezuelan equine encephalitis virus (VEEV), O’nyong’nyong virus (ONNV), coronaviruses (CoV) OC43 and SARS-CoV-2, parainfluenza virus type 3 (PIV3), respiratory syncytial virus (RSV), vesicular stomatitis virus (VSV), and vaccinia virus (VV) (Figure 5A and Figure S2). Several viruses did not completely require SBDS or SPATA5, including influenza A virus (IAV), reovirus T3D (Reo), and herpes simplex virus-1 (HSV1). Ad5, a non-replicating adenovirus vector expressing GFP from a CMV promoter, was largely unaffected by loss of host factors.

**Figure 5.**
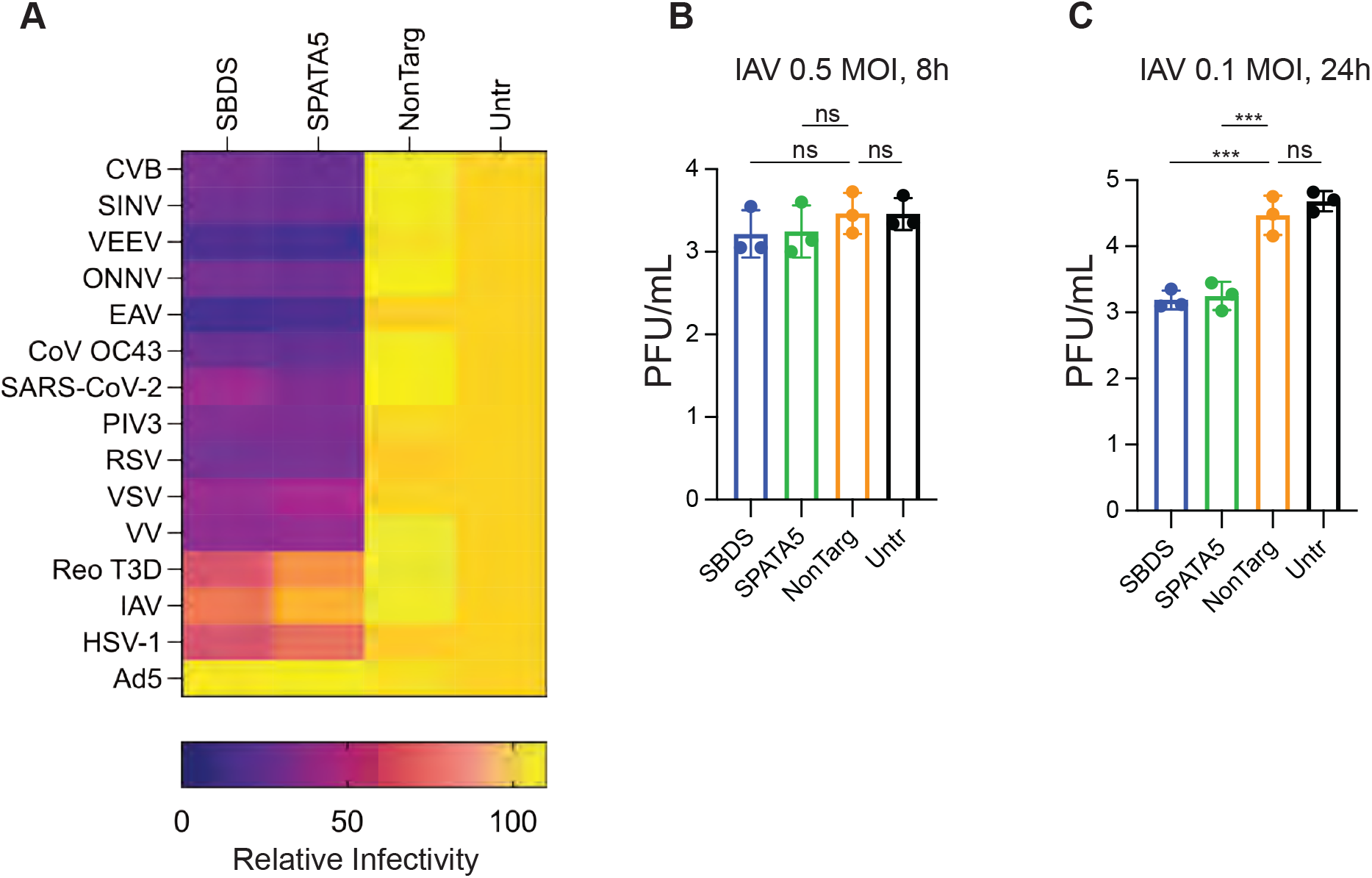
Diverse viruses are susceptible to host factor depletion. A. Viral infectivity heatmap of indicated viruses in control or KO Huh7.5 cells determined by FACS. B-C. Viral titers from supernatants from control or KO Huh7.5 cells infected with 0.5 MOI IAV for 7 hr (B) or 0.1 MOI IAV for 18 hrs (C) determined by plaque assay.

To examine why certain viruses were less affected by loss of SBDS or SPATA5, we used IAV as a model. In the assay presented above, we quantified IAV infectivity by NP staining during a single cycle of replication, and this was not significantly affected by loss of SBDS or SPATA5. We then tested whether dosing and time point factored into the observed phenotype with IAV. We quantified IAV plaque forming units (PFU) in control or KO cells at 0.5 MOI during one round of replication (Figure 5B), or at 0.01 MOI over multiple rounds of replication (Figure 5C). Only in the multi-cycle experiment did we observe reduced IAV titers in SBDS and SPATA5 KO cells relative to control. Similar to HCV replicon data, these results suggest that minor defects in virus replication eventually become amplified in KO cells after several rounds of replication. We speculate that viruses unaffected by loss of SBDS and SPATA5 may produce levels of viral mRNAs that are initially sufficient to outcompete host mRNAs for translational machinery. Once the competitive advantage is lost, by lowering IAV MOI for example, these viruses then become sensitive to host factor depletion.

As loss of SBDS and SPATA5 resulted in reduced replication for most viruses, we hypothesized that depleting these factors may affect cell health. We quantified proliferation rates and found that they were reduced ~5-fold in KO cells as compared to control cells (Figure 6A). This was consistent with visual observations that KO cells grew slowly relative to control cells. These results led us to test whether loss of SBDS and SPATA5 affected gene expression independently of viral replication or in a completely non-viral system. We transduced cells with a non-replicating lentiviral vector expressing the red fluorescent protein TagRFP (SCRPSY-TagRFP). Alternatively, we transfected cells with the lentiviral backbone plasmid (pSCRPSY-TagRFP), or a naked DNA plasmid that expressed TagRFP from a CMV promoter. TagRFP reporter levels were quantified by flow cytometry. In all three cases, TagRFP expression was significantly reduced in SBDS and SPATA5 KO cells relative to control (Figure 6D-F), indicating that the capacity for mRNA translation from viral or non-viral sources was compromised in KO cells.

**Figure 6.**
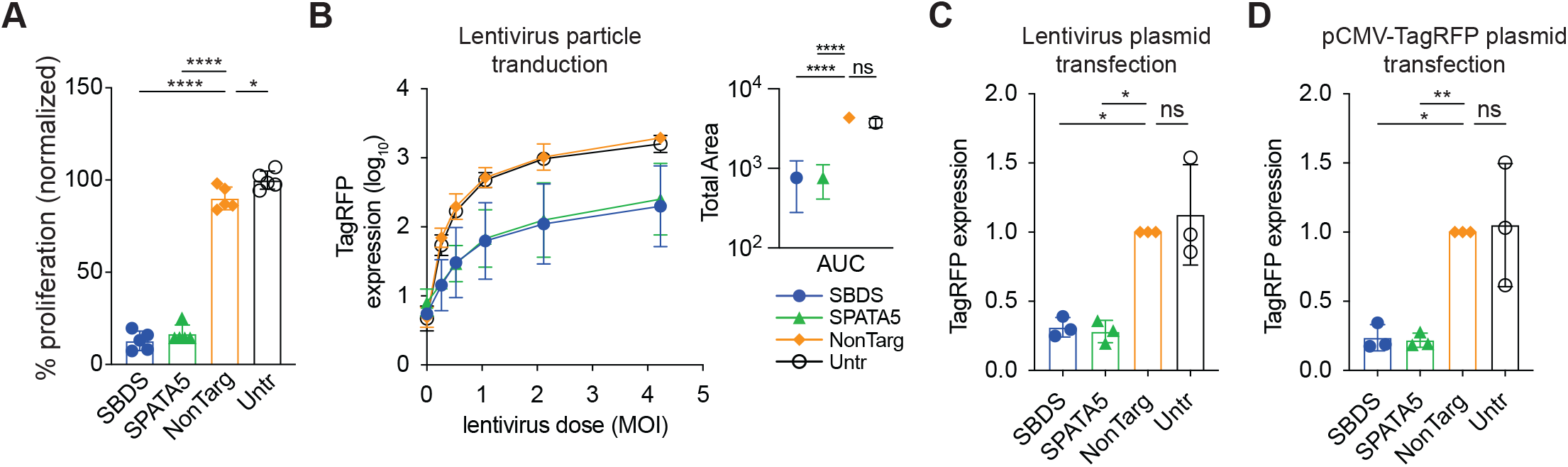
Host factor depletion affect cell health and heterologous reporter gene expression. A. Control or KO Huh7.5 cells seeded overnight were assayed for proliferation by WST-1 assay. Statistical significant was determined by one-way ANOVA, n=5. B-D. FACS-based quantitation of TagRFP expression in KO or control Huh7.5 cells transduced with TagRFP-expressing lentiviral particles (B), transfected with TagRFP-expressing lentiviral plasmid (C), or transfected with a plasmid driving TagRFP gene expression from a CMV promoter. In B, area under the curve (AUC) was calculated from raw data (right panel) and statistical significance was tested by by one-way ANOVA. In C, D, statistical significance was determined by ratio paired t-test of the non-normalized data. n=3, *, P<0.05; **, P<0.01; ***, P<0.001; ****, P<0.0001. Abbreviations: “NonTarg”, non-targeting CRISPR guides; “Untr”, untransduced control.

These global effects on cell health, combined with requirement of these host factors for both viral and non-viral protein synthesis, led us to examine the effect of host factor depletion on cellular ribosome profiles. Lysates of uninfected control and SBDS or SPATA5 KO cells were subjected to sucrose-gradient centrifugation and fractionation. Non-targeting control cells had a polysome profile identical to WT cells (Figure 7A). In contrast, SBDS KO cells had increased 40S and 60S peaks, higher levels of low molecular weight polysomes (LMW), and lower levels of high molecular weight (HMW) polysomes (Figure 7B). Additionally, SBDS KO LMW polysome peaks included “half-mer” peaks that likely contain a 47S pre-initiation complex lacking a joined 60S (Ohtake and Wickner, 1995) and a variable number of ribosomes, suggesting a defect in 60S-joining activity. The SBDS KO profile is consistent with a lack of 60S maturation as the increased 60S peak likely contains pre-60S subunits that are unable to form a competent 80S. Similar to SBDS KO, the SPATA5 KO profile had increased 40S and 60S peaks, reduced 80S, increased LMW polysomes including half-mers, and lower HMW polysomes (Figure 7C).

**Figure 7.**
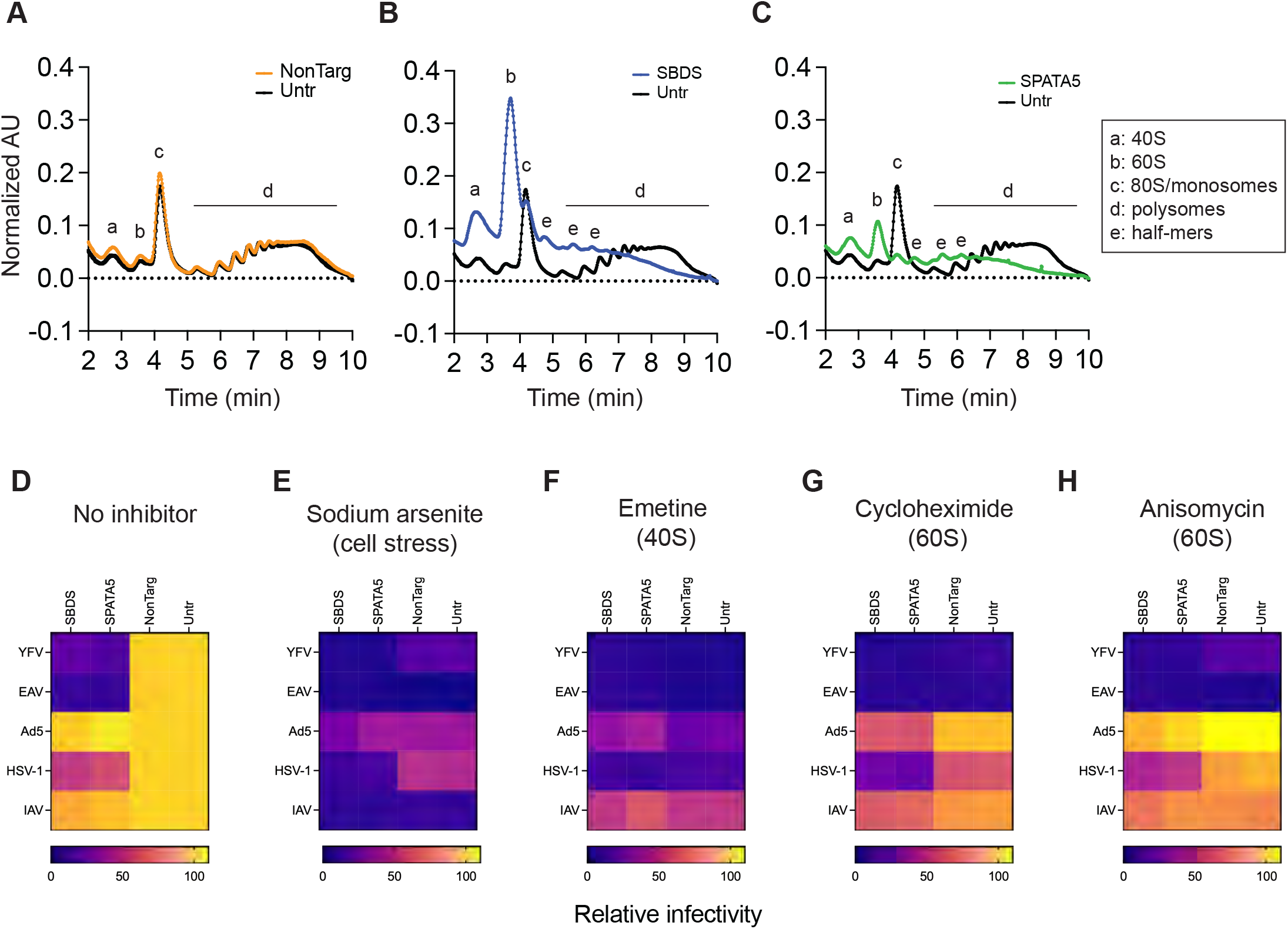
Host factor knock-out cells lack high-molecular weight polysomes. A-C. Sucrose density gradient ribosome fractionation profiles of Non-Targeting control (A), host factor KO cells SBDS (B), or SPATA5 (C), over laid with WT untreated Huh7.5 cell profiles (black). D-H. Viral infectivity heatmaps of control and KO Huh7.5 cells infected with cytoplasmic (YFV, EAV) or nuclear viruses (Ad5, HSV-1, IAV) for 1 hr prior to treatment without (D) or with inhibitors; 50 μM sodium arsenite (E), 2 μM emetine (F), 2 μM cycloheximide (G), or 10 μM anisomycin (H). Infectivity determined by FACS. Abbreviations: “NonTarg” or “NT”, non-targeting CRISPR guides; “Untr”, untransduced control.

The polysome profiles indicate that silencing of SBDS and SPATA5 may result in specific defects in the 60S particle, or its ability to form an 80S monosome. Accordingly, we hypothesized that chemical inhibitors targeting specific stages of translation would phenocopy the viral susceptibility phenotypes of KO cells. Four translation inhibitors with different mechanisms-of-action were chosen; sodium arsenite induces cell stress (Duncan and Hershey, 1987), emetine binds the 40S subunit (Wong et al., 2014), cycloheximide binds the 60S E site, and anisomycin binds the 60S A site (Dmitriev et al., 2020). Conditions for each inhibitor were selected that did not cause host toxicity over a 24h treatment period (data not shown). We then assayed viral replication/gene expression with and without inhibitor treatments 1 hr post-infection for viruses with differing dependencies on SBDS and SPATA5 (YFV, EAV – strong dependency; HSV-1, IAV – moderate dependency; non-replicating Ad5 vector – weak dependency.) Without inhibitor treatment, KO cells were more refractory to YFV and EAV than to Ad5, HSV-1, and IAV, as observed previously (Figure 7D). Cell stress-induced translation inhibition by sodium arsenite reduced replication of all viruses in control and KO cells (Figure 7E). Similarly, blocking translation initiation on the 40S subunit by emetine also inhibited all viruses (Figure 7F). However, inhibition of translation elongation by targeting the 60S subunit with either cycloheximide or anisomycin treatment reduced replication YFV and EAV, both strongly dependent on SBDS and SPATA5, to a much greater extent than the other viruses (Figure 7G-H). The patterns of inhibitor susceptibility correlate with KO cell viral replication phenotypes and support the hypothesis that, under the experimental conditions used here, viruses with strong sensitivities to SBDS and SPATA5 depletion have a unique dependence on the 60S subunit compared viruses that are less affected.

Together, our data suggest that in KO cells, viral protein production can only achieve a certain threshold, or “ceiling”. If that threshold is not surpassed, as it is in wild-type cells, viral replication is suboptimal. This limited protein production correlates with defects in the ribosome profiles of KO cells. One possibility is that the affected viruses require de novo ribosome formation. For example, RPL40-containing ribosomes have been shown to be required for optimal VSV infectivity (Lee et al., 2013). To determine whether de novo ribosome production may be affected, we treated control and KO cells with the alkyne-containing 5-Ethynyl-uridine (5-EU). 5-EU is a uridine mimic that is incorporated into all RNA, but the high abundance of ribosomal RNA (rRNA) relative to other RNAs allows 5-EU to be used a surrogate for potential effects on rRNA production. SBDS and SPATA5 KO cells were 5-EU positive, but they exhibited lower 5-EU incorporation as compared to control cells (Figure 8A), suggesting that de novo production of ribosomal rRNA may be reduced. To determine whether KO cells had defects in the maturation of de novo ribosomes, we assessed rRNA processing by Northern blotting, with probes that detect specific forms of maturing rRNA as ribosomes are formed (Tafforeau et al., 2013). Loss of SBDS and SPATA5 resulted in increased 47S and 34S RNAs, suggesting defects of early cleavage reactions in the 5’ETS (Figure 8B and S4A). Cells lacking SPATA5 additionally accumulated 32S pre-RNA, suggesting impaired ITS2 processing. In yeast, when 60S ribosome biogenesis is disrupted, certain ribosome biogenesis proteins such as Rpl24 are retained with pre-60S particles and thus have defective nuclear shuttling. To determine whether KO cells display altered biogenesis factor localization, we assessed localization of RSL24D1, the human homologue of yeast Rpl24, by immunofluorescence and confocal microscopy. SBDS and SPATA5 KO cells accumulated more RSL24D1 in the cytosol than control cells, suggesting aberrant recycling of RSL24D1 in the absence of these proteins (Figure 8C and S4B). These data complement polysome profiling and inhibitor studies to suggest that SBDS and SPATA5 are involved in the maturation of ribosomes required for optimal viral protein synthesis. The phenotypes of the SPATA5 knockout cells are also consistent with the reported role of the SPATA5-like yeast protein Drg1 in regulating 60S maturation. (Lo et al., 2010). Drg1 has two ATPase domains, and its ATPase activity is required for regulating 60S formation. To test whether the enzymatic activities SPATA5 are required for viral infection, we inactivated ATPase activity by introducing single (glutamic acid/E to glutamine/Q) and double (2x) Walker B mutations in the two ATPase domains (Figure 8D). Wild-type or enzymatic mutants of SPATA5 were expressed in SPATA5 KO cells (Figure S4). In contrast to WT SPATA5, the mutant proteins failed to complement YFV infection (Figure 8E), indicating that that viral infectivity depends on catalytically active SPATA5. Together, our data provide evidence to suggest that human SPATA5 is a functional ortholog of yeast Drg1.

**Figure 8.**
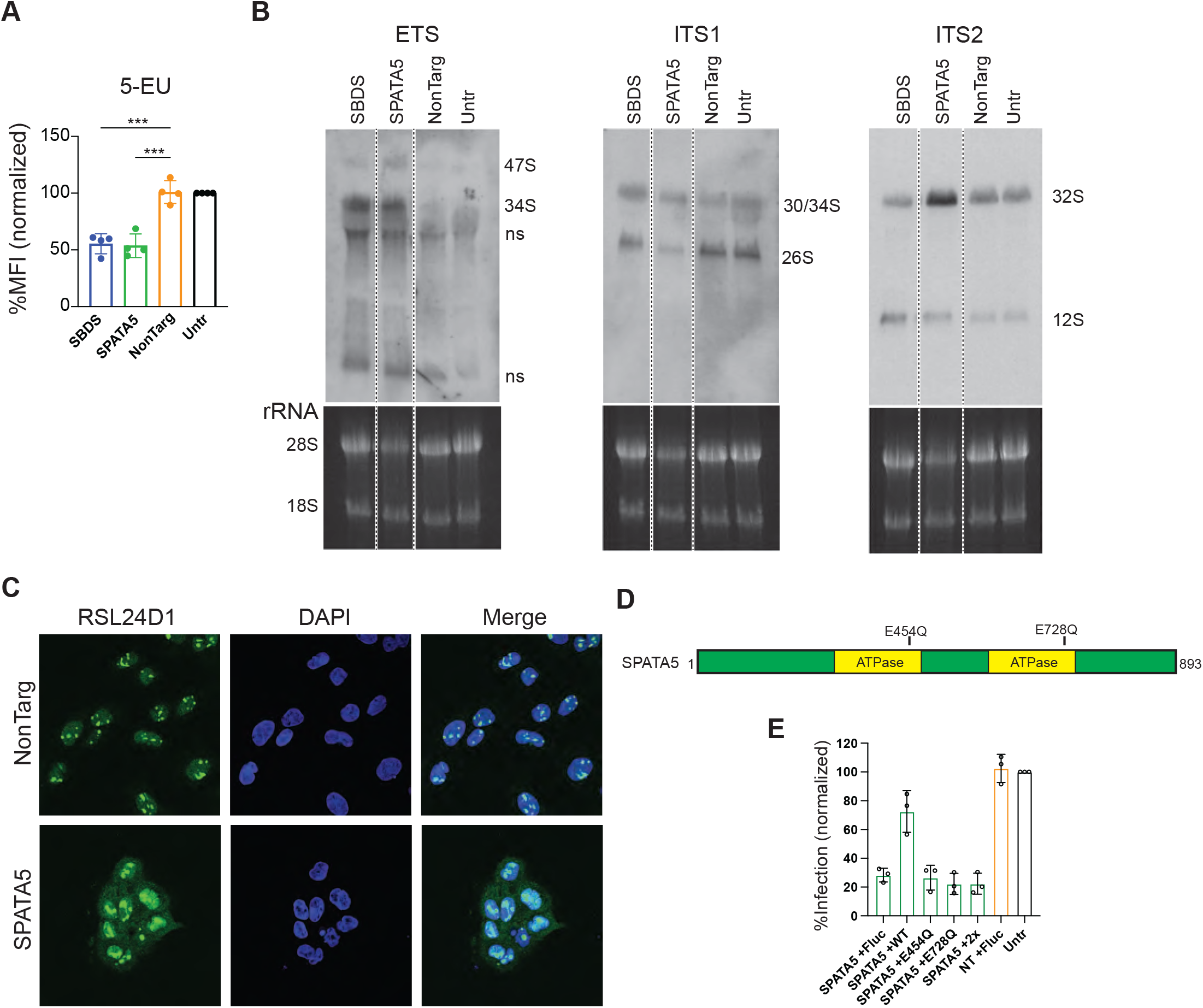
SBDS and SPATA5 and GUK1 regulate ribosome biogenesis. A. Normalized MFI of control or KO Huh7.5 cells treated with 5-EU, labeled by click-chemistry and analyzed by FACS. B. Northern blots of control or KO cell RNA labeled with ETS, ITS1, and ITS2 probes. Images of rRNA gels prior to membrane transfer are shown below blots. Band sizes of rRNA species are labeled. C. Control Non-Targeting or SPATA5 KO Huh7.5 cells were stained with anti-RSL24D1 antibodies (green) and DAPI (blue) and imaged by confocal microscopy. D. SPATA5 protein domain cartoon indicating catalytic residues targeted by site-directed mutagenesis. E. YFV-Venus infectivity assayed by FACS in control or KO cells transduced with lentivirus expressing wildtype or mutant SPATA5 (E), or Fluc control. Abbreviations: “ns”, non-specific; “NonTarg” or “NT”, non-targeting CRISPR guides; “Untr”, untransduced control.

## DISCUSSION

In this study we used a FACS-based negative selection CRISPR-screening approach to identify SBDS and SPATA5 as important regulators of a diverse panel of viruses. We discovered that loss of these host factors impairs ribosome formation and viral protein synthesis, leading to reduced viral replication. Viral sensitivity to host factor loss paralleled phenotypes produced by treatment with translation inhibitors that target the 60S subunit of the ribosome, suggesting that that some viruses may have specific ribosome requirements for replication.

In the CRISPR screen, 19 of the top 93 (<1% FDR cut off) host factor candidates were annotated to 60S ribosome biogenesis factors, and 7 were large subunit (RPL) proteins. In contrast, no 40S biogenesis factors or small subunit (RPS) proteins were found in the top <1% FDR. Expanding the candidate list to the top <5% FDR, a total of 16 RPL and 3 RPS candidates are present, emphasizing the critical role that 60S and RPL proteins may play in flavivirus replication. Indeed, the importance of individual RPL proteins for viral infectivity has been demonstrated for viruses across divergent host species; mammalian RPL40 is important for VSV infection (Lee *et al*., 2013), *Aedes aegypti* RpL23 and RpL27 are required for ZIKV replication in mosquito cells (Shi et al., 2021), and several *Saccharomyces cerevisiae* 60S RPL are required to maintain L-A virus in yeast (Ohtake and Wickner, 1995).

Viruses have evolved numerous strategies to hijack their host’s translational machinery. For example, some viruses use specific RNA structures to form internal ribosome entry sites (IRES) to initiate translation, while other viruses use methods to steal host mRNA components to mimic cellular mRNAs. The viruses that we found to be sensitive to loss of SBDS and SPATA5 do not have any one single translation-hijacking strategy in common. Additionally, not all viruses were affected equally, with non-replicating Ad5 vector being the least sensitive. IAV in a single-cycle assay was also unaffected by loss of SBDS and SPATA5. However, when a lower dose of virus was used in a multi-cycle assay, replication was indeed affected. We speculate that loss of these individual host factors does not completely abolish infectivity, but rather lowers the “ceiling” of viral protein synthesis, thereby imposing an upper limit to optimal viral replication. Viruses that are sensitive to host factor loss could therefore have a lower tolerance for specific perturbations to the host translation machinery. Conversely, the less sensitive viruses may generate high levels of mRNA that outcompete host mRNAs for translational machinery, thereby allowing these viruses to replicate productively, or express genes in the case of the non-replicating Ad5 vector. This model is further supported by demonstrating that TagRFP synthesis from non-replicating lentivirus or even naked plasmid DNA was also dependent on SPATA5 and SBDS, suggesting that the observed translational defects extend beyond viral protein synthesis. Accordingly, cells lacking these two host factors had reduced 5-EU incorporation and proliferation, as would be predicted if ribosomal production and total protein synthesis were impaired, respectively.

Our data suggest a direct link between loss of mature 60S subunits and diminished viral translation. Loss of SBDS in host cells likely produces a pool of pre-60S subunits that are unable to form mature 80S monomers due to lack of EIF6 removal from pre-60S subunits that have been exported from the nucleus (Weis et al., 2015). The AAA ATPase SPATA5 shares homology to yeast Afg2/Drg1, which has been shown to regulate the release and recycling of 60S maturation factors. In both SBDS and SPATA5 KO cells, the presence of half-mers and an increased 60S peak by ribosomal fractionation suggests an accumulation of pre-60S subunits (Figure 5B and 5D), which may outnumber mature, 80S-competent subunits. Notably, in a separate study, SPATA5 was independently discovered along with C1ORF09 to have a similar role in maturation of pre-60S subunits (Ni et al., 2022). C1ORF09 was a statistically significant hit in our CRISPR screen but was not selected for follow-up studies.

In summary, these studies demonstrate that by technically revamping more traditional virus-focused CRISPR screens, previously uncharacterized viral host dependency factors can be identified, revealing new insight into the requirement of specific host proteins for protein synthesis and viral replication. Our data indicate that diverse viruses may have specific requirements for 60S ribosome particles to achieve a threshold of viral protein synthesis that supports optimal viral replication. It is tempting to speculate whether inhibitors of these host factors could lower the “ceiling” of maximal viral protein synthesis, thereby imparting therapeutic antiviral effects. Indeed, a recent study demonstrated this proof of concept by inhibiting SARS-CoV-2 translation with Plitidepsin, which targets the translation factor eEF1A (White et al., 2021).

## MATERIALS AND METHODS

### Cell lines

A549, U-2 OS, HEK293T, MDCK and Huh7.5 cells (from C. Rice, The Rockefeller University) and all derivatives were grown in DMEM (Gibco) supplemented with 10% fetal bovine serum (FBS) and 1X non-essential amino acids (Gibco). BHK-21J cells were grown in MEM supplemented with 10% FBS and 1X non-essential amino acids. L929 cells were grown in DMEM supplemented with 5% FBS, 1% penicillin-streptomycin (Sigma) and 0.1% amphotericin B (Fisher Scientific). All cells were maintained at 37C in 5% CO2. Cells expressing selectable markers were grown in complete media supplemented with puromycin (Sigma) ranging from 0.5-4 μg/mL or blasticidin (Gibco) ranging from 10-20 μg/mL, depending on the cell line. Cell lines were tested for mycoplasma using a PCR assay (Venor GeM Mycoplasma Detection Kit, Sigma).

### Viruses

The generation and production of the following viruses have been previously described (Schoggins et al., 2011), (Schoggins et al., 2012), (Schoggins et al., 2014), (Richardson et al., 2018), (Mar et al., 2018): YFV-17D, YFV-Venus, CVB-GFP, SINV-GFP, DENV-GFP, ZIKV-GFP, HCV-Gluc, PIV3-GFP, RSV-GFP, VSV-GFP, EAV-GFP, VEEV-GFP, VV-GFP, ONNV-GFP, WSN IAV. Reovirus type 3 Dearing (T3D) was provided by TS Dermody and stocks were generated in L929 cells, gradient purified, and quantified by plaque assay in L929 cells as previously described (Kobayashi et al., 2007), (Smith et al., 1969), (Virgin et al., 1988). Ad5-GFP was propagated in HEK-293 cells. HSV1-GFP strain 17 (provided by D. Leib) and SARS-CoV-2 (USA-WA1/2020 BEI #NR-52285) were propagated in VeroE6 cells. Coronavirus OC43 (ATCC VR-1558) was propagated in HCT-8 cells and detected by antibody staining (MAB9012, Millipore).

### Lentivirus pseudoparticle production and transduction

LentiCRISPRv2 (Puromycin, a gift from Feng Zhang, #52961, (Sanjana et al., 2014) and lentiCRISPRv2-Blast (a gift from Mohan Babu, #83480) were purchased from Addgene. Oligos encoding single guide RNA sequences (Table S3) were annealed and ligated into LentiCRISPRv2 vectors linearized with Esp3I according to Addgene protocol. The SCRPSY-TagRFP and SCRBBL lentiviral vector for overexpression have been previously described (Kane et al., 2016; Richardson *et al*., 2018). Infectious lentivirus was produced by co-transfecting plasmids encoding VSV-g, Gag-Pol, and lentivirus vectors at a 2:8:10 μg ratio to HEK293T cells using Xtreme-Gene9 (Roche) and lentivirus-containing supernatants were collected 72hr post-transfection. For generating pools of single-gene knock outs, cells were reverse-transduced by plating cells in the presence of lentivirus expressing two different gene-targeting sgRNAs for 48 hrs, then cells were grown in the presence of selective media for 3-12 days prior to seeding for experiments.

### CRISPR-Cas9 host factor screening

Human Brunello CRISPR knockout pooled library was a gift from David Root and John Doench (Addgene #73178) and amplified according to instructions. Production of library lentivirus and transductions of Huh7.5 cells were performed as previously described (Richardson *et al*., 2018). Huh7.5 cells transduced with 100X coverage of CRISPR library lentivirus were amplified and passaged for two weeks prior to screening; one week in complete DMEM supplemented with 4ug/ml puromycin followed by another week in complete DMEM without antibiotics. The day before infection ~8×10^6^ library cells were seeded per 15cm plate to six plates. Each plate was infected with YFV-venus (MOI 5) in 16 mL of DMEM 1% FBS, and after 3 hrs viral entry 10 mL of complete DMEM was added and the plates were incubated overnight. Twenty-four hours post infection, infected library cells were rinsed with PBS, harvested with trypsin and pooled. Pelleted cells were resuspended in PBS supplemented with 2% FBS and 0.5 mM EDTA, filtered by 100micron filter, and stored on ice before sorting at the Children’s Medical Center Research Institute Flow Cytometry Facility using a FACSAria II (Becton Dickenson) flow cytometer. GFP-negative gated cells were collected in PBS supplemented with 50% FBS and 50 mM HEPES and pelleted. Genomic DNA (gDNA) was extracted from sorted cell pellets as previously described (Richardson *et al*., 2018). Uninfected library cells (3×10^6^) were harvested and gDNA was extracted for controls. The entire screen procedure was performed three separate times, and each time 3.5-5.4×10^7^ cells were sorted by FACS, representing an average of 600-fold library coverage for each screen. Library amplicons of sgRNA sequences in each gDNA sample were generated by PCR using 6-10ug gDNA per 100μL PCR reaction (Doench et al., 2016). Each sample was amplified in 4 parallel 100 μL PCR reactions using barcoded P7 primers (Table S3), then pooled and purified by AMPure XP beads (Agencourt). Prior to sequencing, samples were analyzed by a Bioanalyzer high Sensitivity DNA kit (Agilent) and quantified by qPCR using the KAPA Library Quantification Kit for Illumina. Pooled amplicon library samples were sequenced using Illumina NextSeq 500 with a single end 75bp read configuration. Each sample was subjected to approximately 10-15 million reads. Raw FASTQ files were de-multiplexed and trimmed to extract 20bp sgRNA prior to mapping reads to reference Brunello sgRNA sequences. MAGeCK (Li *et al*., 2014) software was used for data analysis; median normalization was used to adjust for the effect of library sizes and read count distribution. Positively selected sgRNAs and genes were identified with default parameters.

### Viral infection assays

Cells were seeded at 80,000-100,000 cells per well to 24 well plates the day before infection. Virus inoculum (MOI 0.01-1) was added to each well in 200 μL of DMEM 1% FBS for 0.5-3hr, then brought up to 500 μL volume with complete DMEM without or with inhibitor supplemented for the remaining incubation time. At the end of infection time course, media was aspirated, cells were lifted with Accumax (Sigma), mixed with paraformaldehyde (PFA) in PBS to a final concentration of 1% PFA, and fixed for 10 min at room temperature. Fixed GFP-virus infected cells were pelleted and resuspended in PBS supplemented with 3% FBS prior to analysis on a S1000 (Stratadigm) flow cytometer and data were analyzed by FlowJo software. Non-reporter virus infected cells were subjected to fixation and permeabilization (BD Cytofix/Cytoperm #554714), followed by antibody staining: DENV2 and ZIKV, anti-E protein D1-4G2-4-15; CoV OC43, anti-NP Millipore #MAB9013; CoV SARS-CoV-2, anti-NP Sino Biological #40143-MM05; IAV, anti-NP HT103 Kerafast #EMS010; Reovirus, anti-T3D G5 was deposited to the DSHB by Dermody, T.S. (DSHB Hybridoma Product G5). Antibody-stained cells were resuspended in PBS supplemented with 3% FBS and analyzed flow cytometry. For flavivirus viral production experiments, infections were performed as described above, except viral inoculum was removed after 1 hr, cells were washed four times, and supernatants were collected at indicated time points and used for plaque assay or re-infection assay to quantify virus production. HCV-Gluc virus infections were performed with 40,000 cells per well in 48 well plates, HCV-Gluc virus inoculum was removed, cells were washed four times, and supernatants were collected at indicated time points and used for Renilla luciferase assays to quantify viral replication. For SCRPSY-TagRFP lentiviral transduction assays, 50,000 cells were seeded into 48-well plates and lentivirus particles were added to cells in a dose-dependent manner. Cells were collected 24 h later, fixed in 1% PFA, and TagRFP expression was quantified by flow cytometry.

### Transfection studies (replicon, plasmid DNA)

HCV replicons expressing Gaussia luciferase (Bi-Gluc-JFH-SG) were described previously (Schoggins *et al*., 2011). Replicon RNA was transcribed from linearized template plasmids with T7 RiboMAX Express (Promega) and cleaned by MEGAclear (Ambion/ThermoFisher) according to manufacturer instructions. Control and KO cells were seeded at 40,000 per 48 well and transfected with 200 μg RNA in 25 μL OptiMEM using Trans-IT mRNA Transfection Kit (Mirus) according to manufacturer instructions. At each time point, supernatants containing secreted Gluc were collected, and fresh media was replaced until the end of the time course. Gluc Renilla luciferase units (RLU) were measured in 10 μL of each supernatant mixed with 10 μL 2X Renilla lysis buffer (RLB) by Renilla Luciferase Assay Kit (Promega) in white 96 well plates on a Berthold plate reader. For plasmid transfection assays, 200 ng of plasmid pTagRFP-C (Evrogen), which expresses TagRFP from a CMV promoter, or 200 ng pSCRPSY-TagRFP lentiviral backbone plasmid were transfected into 50,000 cells in a 48 well plate using XtremeGene 9 according to manufacturer’s instruction. Cells were collected 24 h later, fixed in 1% PFA, and TagRFP expression was quantified by flow cytometry.

### Viral titer by plaque assay

Infectious YFV-17D particles were quantified by infecting serial 1:10 dilutions of supernatants on BHK-21J cells as previously described (Boys et al., 2020). IAV infectious particles were measured similarly by infecting serial 1:10 dilutions of supernatants on MDCK cells as previously described (Mar *et al*., 2018).

### Viral titer by supernatant reinfection

Infectious DENV2 or ZIKV PRVABC159 particles were quantified by re-infecting 1:5-fold dilutions of supernatants on Huh7.5 cells seeded at 80,000 per well in 24 well plates. Supernatant-infected Huh7.5 cells were lifted with Accumax and fixed in 1% PFA 24 hrs post infection. Infected cells were stained with anti-E protein D1-4G2-4-15 antibody prior to flow cytometry. The percent infected Huh7.5 cells per condition was used to calculate the proportion of infectious particles per volume of supernatant as previously described (Grigorov et al., 2011).

### Guide-resistant and site-directed mutant complementation constructs

SBDS in pShuttle was purchased from GeneCopoeia (Cat# GC-T0315). SPATA5 cDNA was generated from U-2 OS RNA using SuperScript IV First-Strand Synthesis System (Invitrogen) and SPATA5 was amplified from cDNA using ExTaq (Takara) with primers designed in-house (Table S3) that contain BP overhangs and recombined into pENTR using BP Clonase (ThermoFisher). Guide-resistant pENTR or pShuttle constructs were generated by modification of PAM sites by introducing silent mutations using site directed mutagenesis (Table S3). SPATA5 pENTR construct was modified for both guide 1 and guide 2 PAM sites in each “dual” CRISPR KO treatment, whereas SBDS pENTR plasmid was only modified for one PAM site each because one sgRNA PAM site was outside of the protein coding region. Silent site-directed PAM mutations were confirmed by sequencing, and guide-resistant constructs were moved to the blasticidin-selectable pSCRBBL lentivirus expression vector using Gateway LR clonase II (ThermoFisher). Guide-resistant pSCRBBL plasmids were used as templates for site directed mutagenesis using oligos to introduce catalytic mutations (Table S3), and resulting candidate plasmids were sequenced to confirm nucleotide changes. Blasticidin-selectable WT and mutant pSCRBBL vectors were used to generate infectious lentivirus and were transduced to KO cells generated by dual selection of guide 1 and guide 2 sgRNAs in LentiCRISPRv2-puromycin vectors to test complementation.

### Western blots

Cells were lysed using M-PER (ThermoScientific) supplemented with 1x protease inhibitor tablet (Roche) according to manufacturer instructions. Lysates mixed with 2x Lammeli Sample Buffer (BioRad) supplemented with beta-mercaptoethanol were boiled for 5 min and separated on 12% SDS-PAGE gels prior to transfer to methanol-activated PVDF membranes using the Trans-Blot Turbo system (Bio-Rad). Membranes were blocked in Tris-buffered saline with Tween-20 (TBS-T) and 5% dry milk, then probed with dilutions of primary and secondary-HRP antibodies (anti-SBDS Novus #NBP2-22594, anti-SPATA5 Novus #NBP2-38302, anti-actin-HRP Sigma #A3854, anti-mouse-HRP, anti-rabbit-HRP) in TBS-T and developed to X-ray film using Clarity Western ECL reagent (Bio-Rad). Blots re-probed with additional antibodies were first stripped with Re-blot Plus (Millipore #2504) solution and incubated with TBS-T plus milk blocking prior to follow up primary and secondary antibody treatments.

### 5-EU labeling of rRNA

Cells were seeded at 100,000 per well in 24 well plates the day before, then incubated with 500 μM 5-EU (Click Chemistry Tools #1261) diluted in complete media for 2 hrs, rinsed, lifted with Accumax, and fixed in 1% PFA. Following fixation, cells were permeabilized by PBS supplemented with 3% BSA and 0.1% saponin for 15min. 5-EU was labeled in permeabilized cells using the Click & Go Kit (Click Chemistry Tools #1263) with 2 μM Alexa Fluor 488 azide (Click Chemistry tools #1275) according to manufacturer instructions. Labeled cells were washed twice and resuspended with PBS containing 3% FBS prior to analysis by flow cytometry.

### Polysome fractionation

KO or control Huh7.5 cells were seeded at 6.5×10^6^ per 15cm plate 24 hrs prior to experiment. Uninfected control or KO cells were lifted with trypsin and lysed in polysome lysis buffer (20mM Tris-HCl pH 7.4, 5mM MgCl_2_, 100mM NaCl, 1X protease inhibitor, 0.15% NP-40, 100ug/ml cycloheximide, 1:400 RNasin). For infected cell polysome analysis, 15cm plates of WT Huh7.5 cells were infected with ~MOI 5 YFV-17D or YFV-17D-venus for 24hr prior to harvest. Infected cells were harvested and treated similarly, except the concentration of NP-40 in polysome lysis buffer was increased to 0.5% to inactivate virus. Cell lysates were clarified by centrifugation at 12,000xg for 10 min at 4C and layered on top of discontinuous gradients containing layers of 60%-0% sucrose in 14×95mm ultracentrifugation tubes. Gradients were centrifuged at 150,000xg for 3 hr at 4C in a Sw40ti rotor. Separated gradients were analyzed by A254 with a BioRad BioLogic LP and 500 μL fractions were collected with a BioRad BioFrac.

### rRNA Northern blotting

RNA from 5×10^6^ cells was extracted using 1 mL TRIzol according to manufacturer instructions. 10 μg of purified RNA was analyzed for rRNA processing by Northern blot using NorthernMax kit (Invitrogen) and probes listed in Table S3.

### Immunofluorescence for RSL24D1

Control or KO cells were seeded to poly-lysine treated 12mm glass coverslips in 24 well plates prior to fixation with 4% paraformaldehyde. Fixed cells were permeabilized with PBS plus 0.1% Triton X-100 for 10 min and blocked for 1 h with PBS containing 5% BSA and 0.1% Tween-20. Coverslips were incubated with rabbit anti-RSL24D1 (ProteinTech) diluted 1:100 in PBS with 1% BSA and 0.3% Triton X-100 overnight at 4C, counterstained with donkey anti-rabbit Alexa Fluor 488 (Invitrogen), and mounted to slides with Vectashield containing DAPI. Cells were imaged by a Nikon confocal microscope and analyzed by FIJI.

## ACKNOWLEDGEMENTS

We thank Neil Alto, Julie Pfeiffer, Terence Dermody and Charles Rice for kindly sharing reagents. We gratefully acknowledge technical assistance provided by the Children’s Medical Center Research Institute Flow Cytometry Facility and the McDermott Center Next Generation Sequencing Core. This work was supported in part by the Clayton Foundation for Research (J.W.S.) and by NIH grants AI117922, AI158125 (J.W.S.) and GM125812 (M.B.).

## AUTHOR CONTRIBUTIONS

M.B.O. and J.W.S designed the project. M.B.O., J.L.E, S.J., and C.N. performed the experiments. A.K. and C.X. processed the CRISPR datasets. All authors contributed to data analysis and interpretation. M.B.O and J.W.S. drafted the manuscript text, and all authors contributed to editing the manuscript.

## SUPPLEMENTAL FIGURE LEGENDS

**Figure S1.**
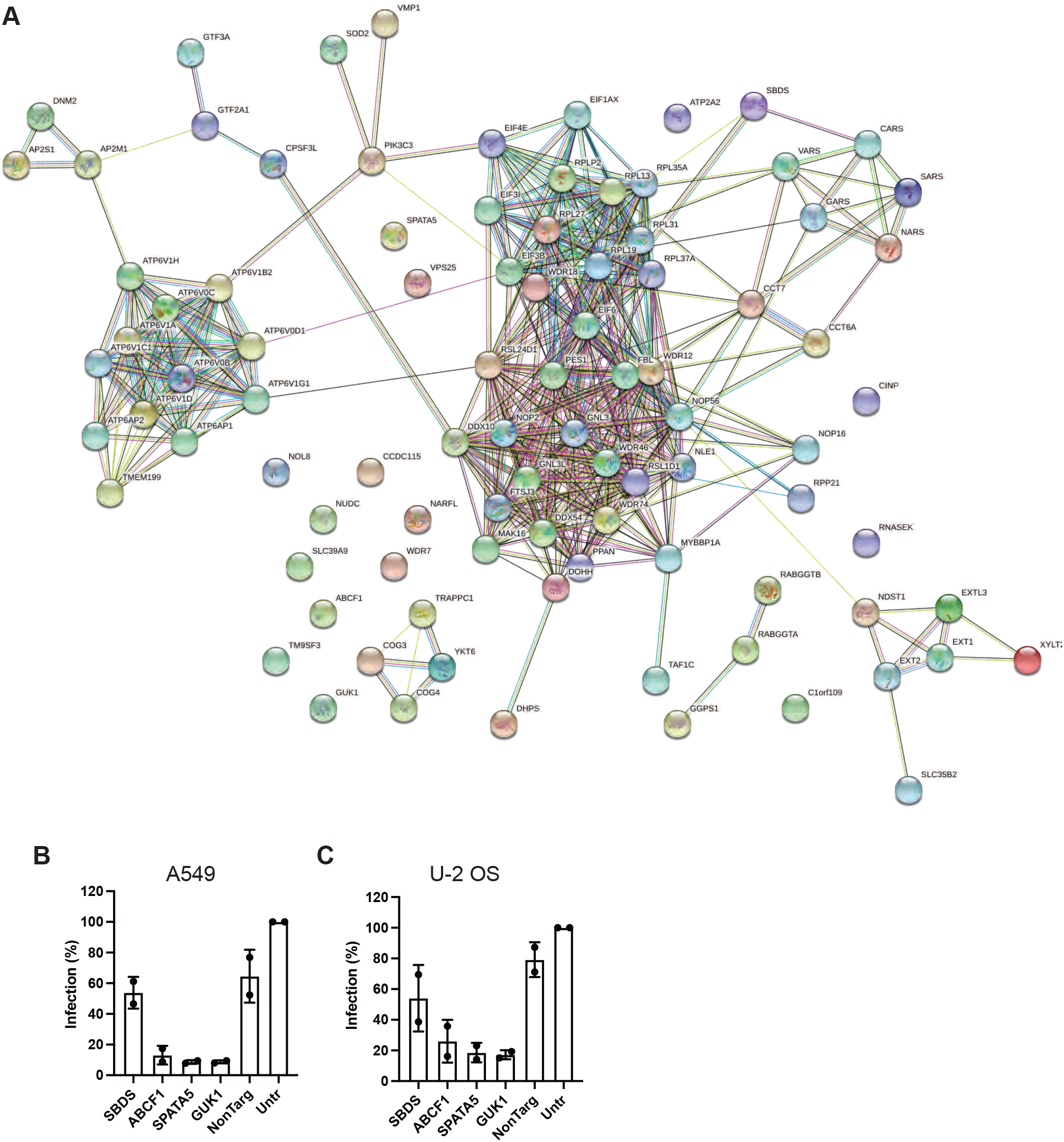
60S biogenesis factors identified in CRISPR screen cluster by STRING. STRING network plot of <1% FDR CRISPR screen results generated by https://string-db.org. B-C. YFV-Venus infectivity assayed by FACS in A549 (B) or U-2 OS (C) control and host factor KO cells.

**Figure S2.**
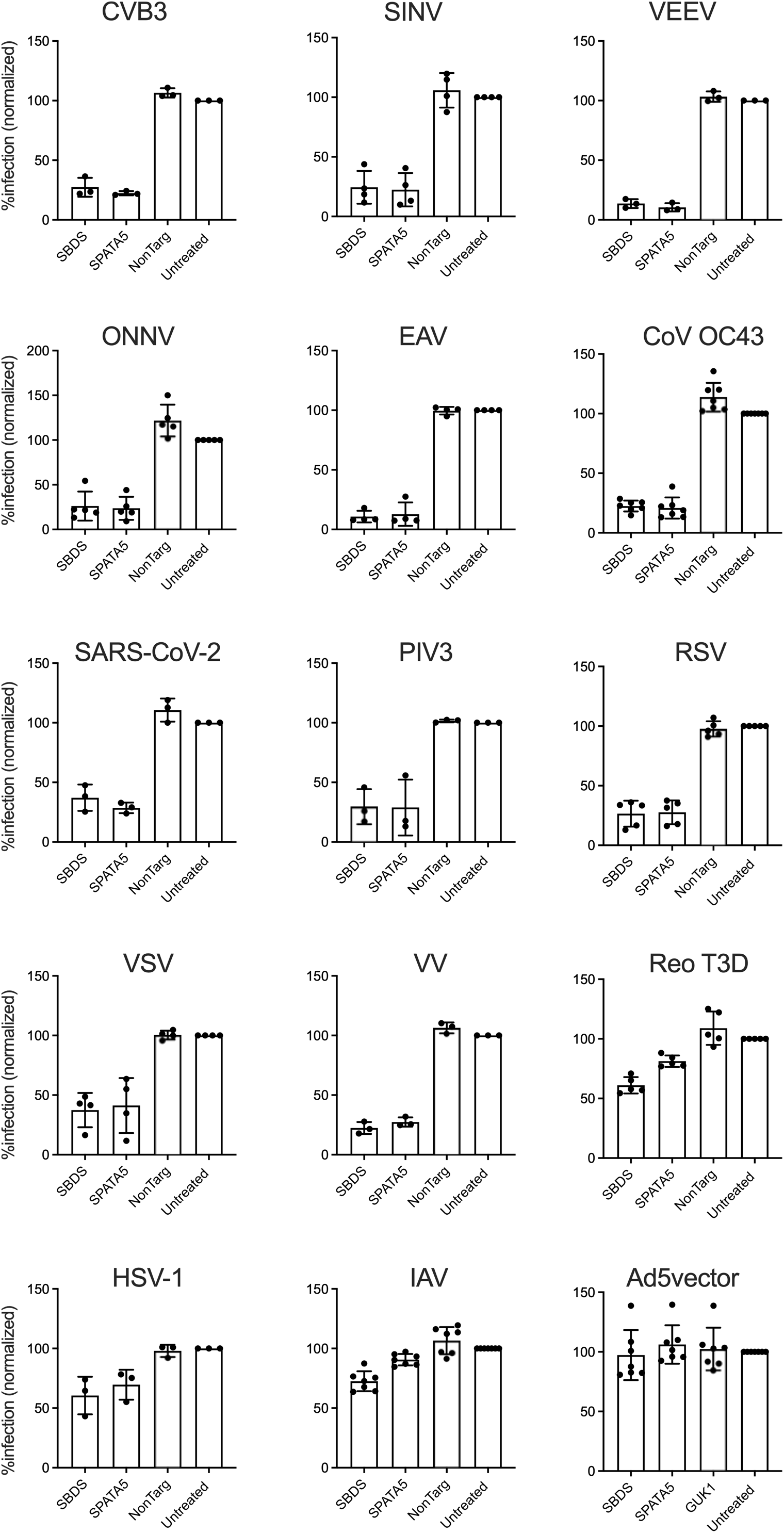
Diverse viruses are susceptible to loss of SBDS, ABCF1, SPATA5 and GUK1. Viral infectivity of indicated viruses in control or KO Huh7.5 cells determined by FACS measuring GFP or antigen staining.

**Figure S3.**
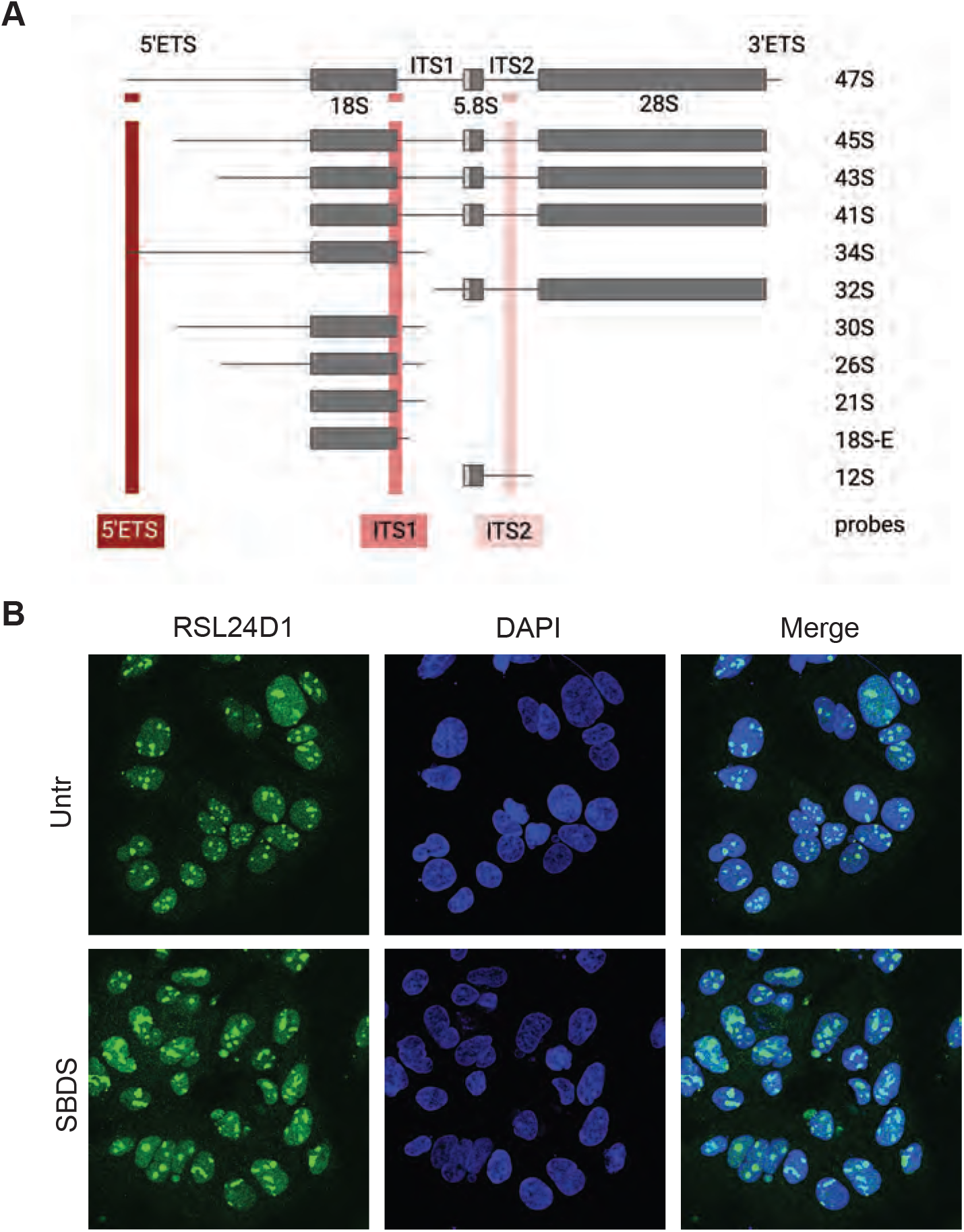
Host factor KO cells have altered RSL24D1. A. Schematic of Northern blot probe binding sites in rRNA. B. Control Untr WT or SBDS KO Huh7.5 cells were stained with anti-RSL24D1 antibody (green) and DAPI (blue), and imaged by confocal microscopy.

**Figure S4.**
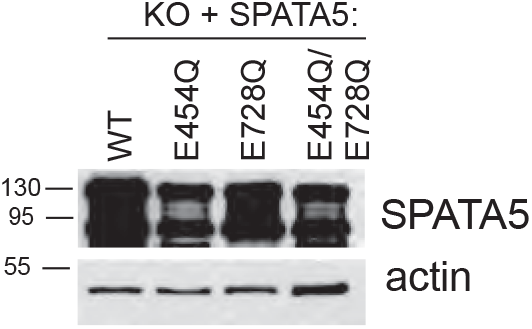
Expression of guide-resistant SPATA5 in SPATA5 KO cells. Lysates of Huh7.5 SPATA5 KO cells complemented with lentivirus-expressed WT or catalytic mutant guide-resistant SPATA5 were analyzed by western blot using anti-SPATA5 and anti-actin antibodies.

