## Supplementary material for "Genome-scale CRISPR screening reveals host factors required for ribosome formation and viral replication": Table S2

Table S2. Viruses tested for replication in Huh7.5 host factor knockout cells

| **Virus name (abbreviation)** | **Family** | **Genome** |
| --- | --- | --- |
| Yellow fever virus (YFV) | Flaviviridae | +ssRNA |
| Zika virus (ZIKV) | Flaviviridae | +ssRNA |
| Dengue virus (DENV) | Flaviviridae | +ssRNA |
| West Nile virus (WNV) | Flaviviridae | +ssRNA |
| Hepatitis C virus (HCV) | Flaviviridae | +ssRNA |
| Coxsakie virus B (CVB) | Picornaviridae | +ssRNA |
| Equine arteritis virus (EAV) | Artirivirus | +ssRNA |
| Sindbis virus (SINV) | Togaviridae | +ssRNA |
| Venezuelan equine encephalitis virus (VEEV) | Togaviridae | +ssRNA |
| O’nyong’nyong virus (ONNV) | Togaviridae | +ssRNA |
| Coronavirus (CoV OC43) | Coronaviridae | +ssRNA |
| Coronavirus (SARS-CoV-2) | Coronaviridae | +ssRNA |
| Parainfluenza virus 3 (PIV3) | Paramyxoviridae | -ssRNA |
| Respiratory syncytial virus (RSV) | Paramyxoviridae | -ssRNA |
| Vesicular stomatitis virus (VSV) | Rhabdoviridae | -ssRNA |
| Reovirus | Reoviridae | dsRNA |
| Influenza A (FluA) | Orthomyxoviridae | -ssRNA |
| Adenovirus 5 (Ad5) | Adenoviridae | dsDNA |
| Herpes simplex virus 1 (HSV-1) | Herpesviridae | dsDNA |
| Vaccinia virus (VV) | Poxviridae | dsDNA |
